# Melanoregulin Deficiency Affects Bone Maintenance and Lean Body Mass

**DOI:** 10.64898/2025.12.14.691656

**Authors:** M.L. Musskopf, V. De P. Gonçalves, S. Tuin, S.W. Wong, K. Boesze-Battaglia, P.A. Miguez

## Abstract

**Objectives:** Studies documented the association of melanoregulin (MREG), a cargo-sorting protein, with its binding partner, the autophagic protein, microtubule-associated protein 1 light chain 3B (LC3B) in macrophages which could affect bone physiology due to the importance of autophagy in osteoclast function. Herein we propose to test the hypothesis that MREG modulates bone remodeling. Therefore, we analyzed the *Mreg*^*dsu/dsu*^ mutant mice for bone mass, growth plate microarchitecture, and bone marrow-derived osteoclast function to understand how lack of MREG affects bone and mass at two different time points.

**Methods:** Mice femurs from wild type and MREG^-/-^ male mice (on C57BL6)/J background) were harvested at 4 and 10 months and imaged by microcomputed tomography to assess bone mass parameters. Femurs were processed for histology by H&E and TRAP staining for assessment of osteoclast numbers. Primary bone marrow-derived macrophages from 3-week-old mice were harvested to assess osteoclast differentiation and function via TRAP, resorptive assay and Western Blot for osteoclast differentiation markers. In addition, a separate cohort of mice were analyzed via EchoMRI to characterize total lean vs. fat whole body mass.

**Results:** There was a statistically significant difference in bone volume of 10-month old mice in wild type vs. MREG^-/-^ with MREG mutation suggesting a preservative effect on phenotypical bone parameters as the mice age. A reduction in adipose tissue but an increase in osteoclast numbers was found histologically in MREG mutant femurs. Bone marrow-derived cells, however, showed reduced osteoclastic function in MREG^-/-^. The mutant mice presented a total lean mass significantly increased compared to wild type per EchoMRI.

**Conclusions:** MREG deficiency seems to impact osteoclast numbers *in vivo* but not *in vitro*, although *in vitro* function was reduced. MREG deficiency favors lean mass preservation over fat accumulation in bones and body composition as mice age. This study provides the foundation for a more in-depth investigation of MREG’s role in bone and systemic metabolism. It is possible that MREG can be a future target for new therapeutic modalities in inflammatory and metabolic bone diseases.

## INTRODUCTION

Bone is a metabolically active tissue with a network structure constituted of different types of cells and regulated by various biological, biochemical and physical factors. The maintenance of a stable bone metabolism requires the continuous differentiation and maturation of different progenitor cells, the preservation of the mineralizing function of osteoblasts, the resorptive capacity of osteoclasts, the secretion of factors that regulate interactions between cells and the extracellular matrix (1-5). Bone remodeling is essential for the formation and maintenance of bone morphology and the regeneration of damaged bone (6, 7), and a dysfunction of any cell type involved in this process can lead to the failure of these preocesses followed by the development of conditions such as osteoporosis (7, 8).

Autophagy plays an important role in maintaining cellular function and it is a complex dynamic process involved in the degradation of intracellular proteins and organelles under different physiological or pathological conditions(1). It is a vital function to maintain cell homeostasis in response to starvation, hypoxia, infection, and other stressful stimuli (1, 3, 7, 9). In bone tissue, autophagy plays key functions in bone marrow mesenchymal stem cells (BMSCs) and derived cells including chondrocytes, osteoblasts and osteocytes (3, 10).

Osteoblasts are specialized mesenchymal-derived cells involved in bone formation, and impaired autophagy in these cells has been shown to lead to decreased bone mass, while inhibiting autophagy in osteocytes results in bone tissue senescence (11-13). Previous evidence showed that altered autophagy inhibits the function of endogenous BMSCs and further promotes the development of osteoporosis (14, 15). *In vitro* studies demonstrated that downregulation/ablation of autophagy markers resulted in the dysfunction or inhibition of osteoblast differentiation (11, 16, 17). Complementary, *in vivo* studies have showed that knockout of autophagy-related genes in mice, resulted in impaired mineralization and reduced bone mass (16, 18).

Moreover, it has been shown that undifferentiated mesenchymal stem cell (MSCs) contain more autophagosomes than differentiated ones, suggesting that autophagy is correlated with MSC differentiation (3). When those cells are exposed to osteogenic stimulation, the accumulated autophagosomes can degrade rapidly, providing the corresponding energy and metabolic precursors for the morphological, function, and metabolic changes required for cell differentiation (19). In corroboration, it is known that during early-stage osteogenic differentiation, the expression of autophagosome marker LC3-II is reduced within 12h, suggesting that these accumulated autophagic vacuoles may serve as a source of rapidly produced energy substrates to support differentiation (20, 21). Frost et al. reported how *P. Gingivalis’* lipopolysaccharide (LPS)1690 affects autophagy assessing LC3-dependent and MREG-dependent processes in green fluorescent protein -LC3-expressing Saos-2 cells (pre-osteoblastic cells) (22). The authors observed that LPS stimulated the formation of very large LC3-positive vacuoles and MREG puncta, as they work together in autophagosome formation.

Further, autophagic proteins are also necessary for osteoclast-directed bone resorption (23, 24) and previous studies have reported that increased authopagy in osteoclasts stimulated cell differentiation and osteoclastogenesis leading to increased bone resorption. I*n vitro* the knockdown of autophagy-related genes in osteoclasts has been shown to significantly reduce the depth and volume of bone resorptive pits (24-28). To further elucidate the role of MREG in bone, we investigated the phenotype of MREG-deficient mice in skeletal bone. Given MREG’s documented role in endolysosomal trafficking and LC3-dependent processes in other cell types, and its expression in bone tissue, investigating its function in bone remodeling is warranted (21, 22). We hypothesize that since MREG is involved in the autophagy process, its deficiency may be implicated in bone physiology and will lead to positive associated bone changes (bone preservation) with maturation of bones. The purpose of this study was to characterize the femoral bone phenotype of the MREG knockout mice and acquire further insight into MREG function not only in osteoclast cell differentiation and function but also evaluate the resulting bone phenotype in the context of development and maturation.

## MATERIAL AND METHODS

### Animals

Melanoregulin (MREG) is the product of the *Mreg*^*dsu*^ gene locus [previously known as dilute suppressor (*dsu*) (29). *Mreg*^*dsu/dsu*^ mice carry the *Mreg*^*dsu*^ allele (mouse accession number Q6NV65), in which the deletion of the first two exons results in an effective null allele. *Mreg*^*dsu/dsu*^ mice (on C57BL6/J genetic background) used in these studies were originally maintained and propagated at the NCI, National Institutes of Health, [generous gifts from Drs. N. Jenkins and N. Copeland (Texas Medical Center)]. Both *Mreg*^*dsu/dsu*^ and MREG^+/+^ [C57Bl6/J mice, (obtained from the Jackson Laboratory)] mice were housed under standard cyclic light conditions: 12-h light/12-h dark and fed ad libitum.

The animal experiment protocol was approved by the Institutional Animal Care and Use Committee (IACUC) at the University of North Carolina at Chapel Hill (IACUC ID: 22-053). The procedures were also conducted in compliance with ethical standards that fully comply with Animal Research: Reporting of *In Vivo* Experiments (ARRIVE) guidelines. Male MREG^+/+^ and MREG^-/-^ mice (on C57BL6)/J genetic background) at 6 weeks of age (20–27 g) were kept in an environment with controlled temperature, humidity, and light cycles, and fed with water and feed ad libitum. Mice were aged up to 10 months. Briefly, mice femurs were harvested at 4 and 10 months and imaged by microcomputed tomography to assess bone mass parameters, femurs processed for histology and stained by hematoxylin and eosin (H&E) for structural and tartrate resistant acid phosphatase (TRAP) for osteoclast numbers assessment. Mice were also scanned via magnetic ressonance to characterize total lean vs. fat mass. Primary bone marrow-derived macrophages from 3-week-old mice were harvested to assess osteoclast function as described below.

### Microcomputed Tomography

After mice reached 4 and 10 months of age, animals were sacrificed under humane conditions as approved by IACUC. Femurs and heads were harvested from 4-5 mice from each group, fixed in 10% buffered formalin for 3 days under constant agitation and stored in PBS at 4°C until dissected for imaging. Sample size was determined from previous studies showing similar bone volume effects (30). The femurs were scanned using a microcomputed tomography (µCT) system (Skyscan 1275, Bruker) at 12.45 µm, 70 kVp, 142 µA using a 0.5 mm aluminum attenuation filter. The three-dimensional images were reconstructed using the software NRecon 1.6.9.8 (SkyScan), Data Viewer 1.5.0 (SkyScan) and CTAnalyzer (CTAn—2003-11SkyScan, 2012 Bruker MicroCT, 1.13.11.0 version) and were used for image re-orientation and analysis, respectively. For the analysis, a standardized region of interest (ROI) included a set of specific number of slices below the growth plate after a set anatomical marker for trabecular bone and cortical bone analysis (50 and 400 slices below growth plate, respectively). The bone microarchitecture parameters evaluated were trabecular and cortical bone volume fraction (BV/TV, %), trabecular thickness (Tb.Th), trabecular number (Tb.N), trabecular separation (Tb.S), as the recommended parameters reported by Bouxsein et al. and as we previously reported (30, 31).

### Histological analysis

After the femurs were demineralized in EDTA ph 7.4 for 4 weeks, they were embedded, non-serial histological sections (6 μm) were obtained and stained by H&E for a qualitative gross evaluation.

For TRAP staining, slides were deparaffinized and rehydrated through graded ethanol to distilled water, and then immersed in TRAP staining solution mix (at 37°C for 30 minutes). The sections were rinsed in distilled water and counterstained with 0.02% fast green for 30 seconds. For quantification, TRAP-positive cells stained in red and containing three or more nuclei on the bone surface, were considered osteoclasts, and quantified in the region of interest immediately below the growth plate as we previously reported (24).

### WT and MREG^-/-^ primary osteoclasts harvest, differentiation and function

In a separate cohort of animals, after euthanasia by CO2, tibias and femurs were harvested from 3 week-old MREG^+/+^ /MREG^-/-^ mice (on C57BL6)/J genetic background). Three independent experiments were performed, each using bone marrow pooled from 3 mice per genotype with triplicates. Bone marrow cells were flushed into the phenol-free α-MEM medium, supplemented with 10% FBS, L-glutamine, nonessential amino acids, and penicillin/streptomycin. Non-adherent cells were harvested after 24 h and re-plated at a density of 1.5x10^5^ cells/cm2 with 30 ng/mL M-CSF (R&D Systems, Minneapolis, MN, USA). After 2 days, the medium was replenished with 30 ng/mL M-CSF and 10 ng/mL RANKL (R&D Systems Minneapolis, MN, USA) for osteoclast differentiation. After 5 days of culture, cells were fixed and stained with TRAP. Equal numbers of bone marrow osteoclast precursors were plated and differentiated on Osteo Assay Surface plates (Corning Lifesciences, Tewksburg, MA, USA) for resorption assay. At day 5, the plate was bleached, and areas of resorption pits were quantified using NIH ImageJ (24, 32).

Bone marrow osteoclast precursors were plated and differentiated by 3 days for Western Blot (WB) analysis. On days 0, 1, and 2 of the assay, the supernatant of each well plate was aspirated and cells harvested. Briefly, RIPA Buffer solution (ThermoFisher Scientific, Waltham, MA, USA) containing protease inhibitors was added into the plates and using a scraper, the cells were detached and transferred to Eppendorf tubes and kept at −80 °C. Protein quantification was performed using the BCA Protein Kit (Pierce BCA Protein Kit, Thermo Fisher Scientific, Waltham, MA, USA) according to the manufacturer’s guidelines and samples were prepared WB to evaluate the protein expression of c-Fos (Cat. #4384s, Cell Signaling, Danvers, MA, USA), NFATc1 (Cat. #sc-7294, Santa Cruz Biotechnology, Dallas, TX, USA) and as a control, β-actin 1 (Cat. #sc-1616, Santa Cruz Biotechnology). In total, 5-10 µg of the total protein lysate was resolved using Criterion TGX precast gel (Biorad, Hercules, CA, USA) and transferred to a nitrocellulose membrane using the Trans-Blot Turbo Transfer System (Biorad) and immunodetected using appropriate primary and peroxidase-coupled secondary antibodies (Goat-Anti-Rabbit IgG, ThermoFisher #31460). Proteins were visualized using enhanced chemiluminescence (ECL, Amersham Bioscience, Chicago, IL, USA). The protein levels were evaluated relative to β-actin via use of Image J (NIH, Bethesda, MD, USA) (24).

### Echo MRI

Metabolic studies including echo-magnetic resonance imaging (EchoMRI) experiments were conducted as previously described (33). Briefly, 7 live mice, WT and MREG mutants, at 10 months of age, were placed in a specially sized, clear plastic holder without sedation or anesthesia. The tube is 2-inches in diameter with 2-5 inches adjustable length and can fit mice up to 100g in weight. There was a total of eight quarter-inch diameter air holes on each side of the tube to ensure air exchange. The tube (with a mouse inside) was then inserted into the tubular space of the EchoMRI™ system. The scans took place for 2 minutes. The fat/lean mass and body water measurements were recorded in grams. No pain or any physical damage was incurred during the measurements.

### Statistical analyses

For all *in vitro* data, three independent experiments were performed, and each experiment was conducted in triplicate sets. Statistical analysis treated the experiment of cells harvested from mice as the unit of analysis (n=3). For µCT, n=5 for WT, n=4 MREG^-/-^ at 4 months; n=5 per group at 10 months. For Echo MRI, the n=7. Histological quantification of osteoclasts: n=4 per group with 3 sections per animal. For both, *in vitro* and *in vivo* assays, data were submitted to the Kolmogorov– Smirnov or Shapiro–Wilk tests to assess homogeneity and data distribution. Student’s *T* test or analysis of variance (ANOVA) were used to determine the differences among groups. All analyses were performed using GraphPad Prism 8 software (GraphPad Software Inc., San Diego, CA, USA). All tests were applied with a 95% confidence level (*p* < 0.05). Data were expressed as mean ± standard deviation.

## RESULTS

### MREG deletion promoted an age-dependent bone-sparing effect on femoral bone

Trabecular structure using µCT showed that there was no difference in trabecular separation (Tr.Sp), number (Tr.N) and thickness (Tr.Th) between MREG^-/-^ and WT mice at 4 months [Figure (fig) 1A, B]. At 10 months, bone parameters were significantly improved for MREG mutants such as Tr.N and bone volume variables (*p*< 0.05). (Fig. 1C, D). Cortical parameters were not statistically different between age groups (Fig. 1E).

**Fig 1.**
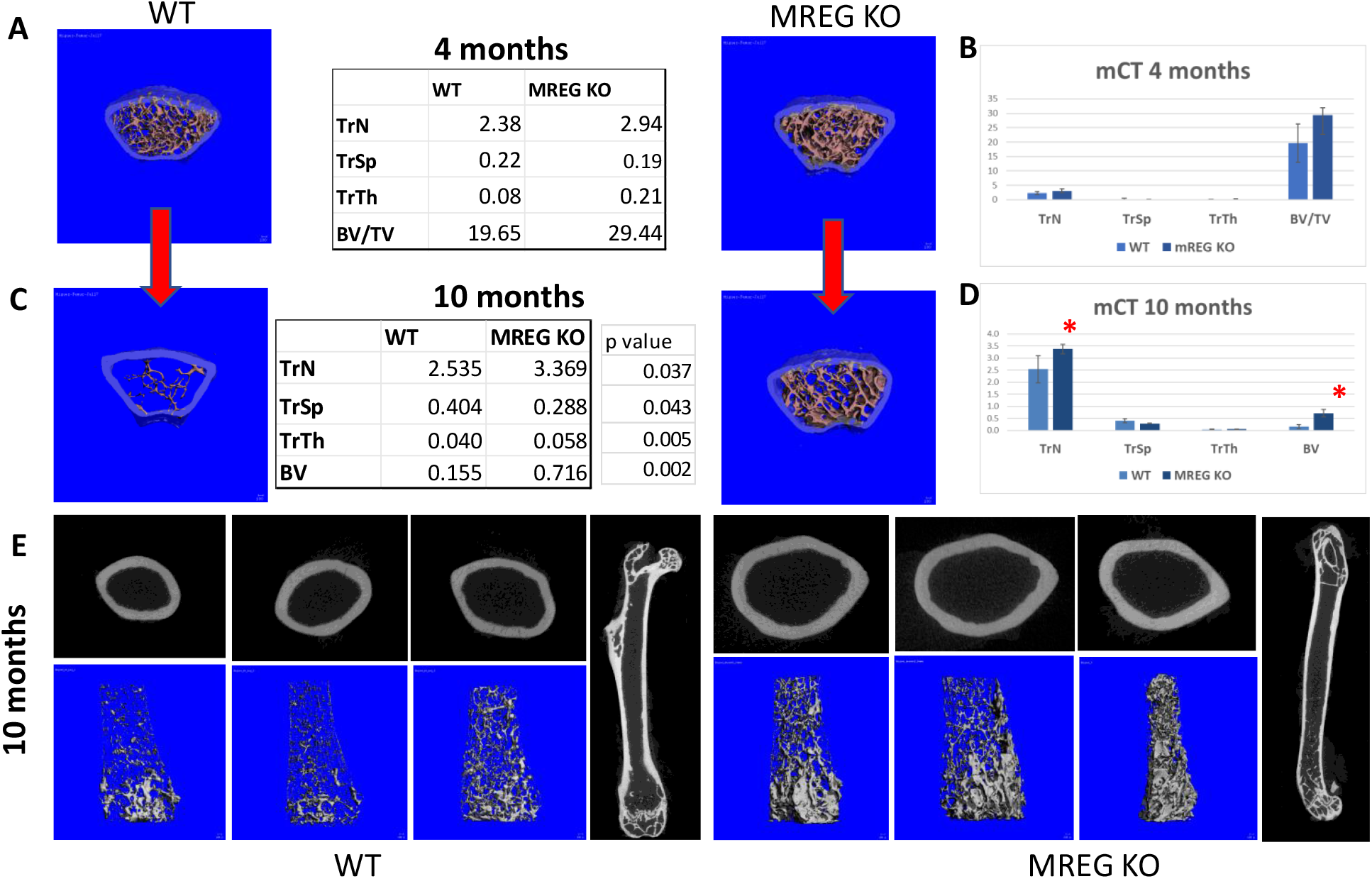
Melanoregulin (MREG) mutation and bone homeostasis at 4 months (A, B) and 10 months (C, D). Trabecular bone and bone volume assessed via microcomputed tomography (mCT) show significant changes, p<0.05, at 10 months of age (C, D). * indicates statistical difference compared to wild-type (WT). At 10 months, cortical parameters were not statistically different (E). 3D rendering of trabecular bone shows clear evidence of improved trabecular fill in MREG knock out (KO) mice at 10 months of age (lower panel). n= 4-5 male mice.

### MREG^-/-^ femurs showed increased osteoclast numbers and decreased evidence of adipocyte population

Qualitative histological assessment of the femurs for gross differences in bone morphometry revealed that marrow-associated adipose tissue was markedly different between MREG^-/-^ and WT. The adipose cells just below the growth plate area was absent in MREG^-/-^ (Fig. 2A, B). Quantititative analysis of osteclasts by TRAP+ multinucleated cell numbers was similar for 4-month-old WT mice (Fig. 3A, B) and MREG^-/-^ (*p* > 0.05). However, in 10-month-old MREG^-/-^ mice there was an increase in osteoclast count (Fig. 3C, D, E) (*p* < 0.05) showing that MREG mutation affected the numbers of osteoclasts as the mice aged.

**Fig 2.**
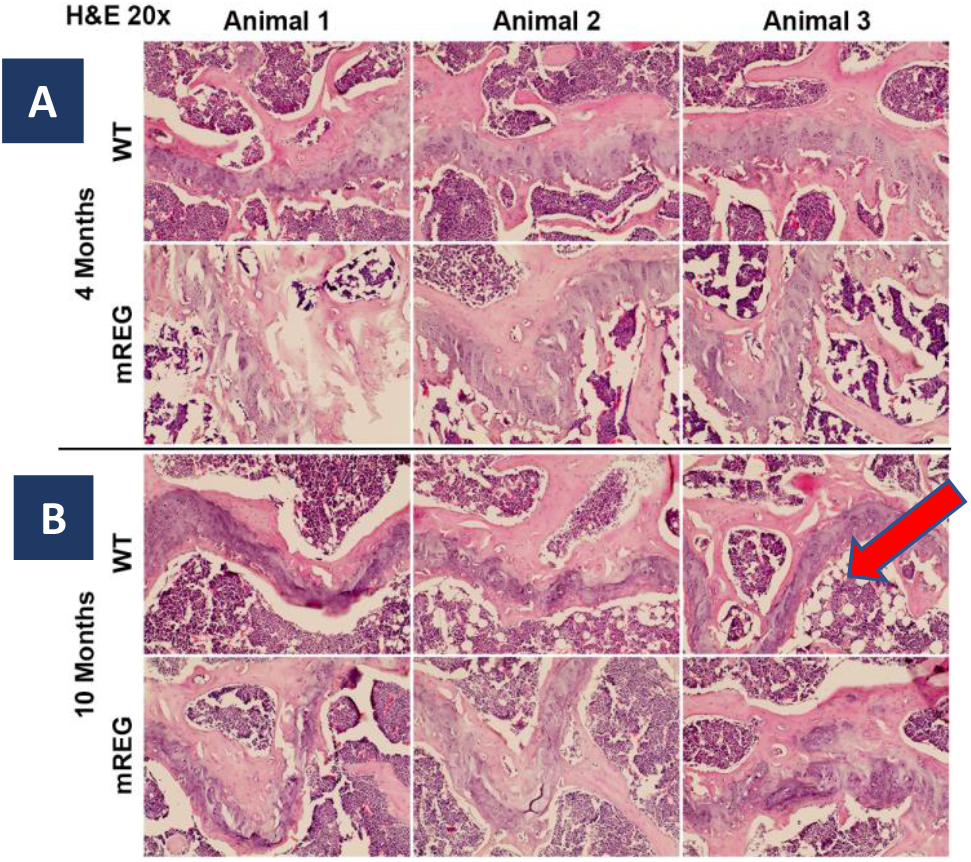
Mouse femurs of MREG mutant mice at 4 months have similar gross morphometric characteristics within the growth plate (A). At 10 months of age, MREG mice showed clear differences in cell composition. Adipocytes are present only in WT mice (red arrow) (B), suggesting a cell fate effect by MREG absence.

**Fig 3.**
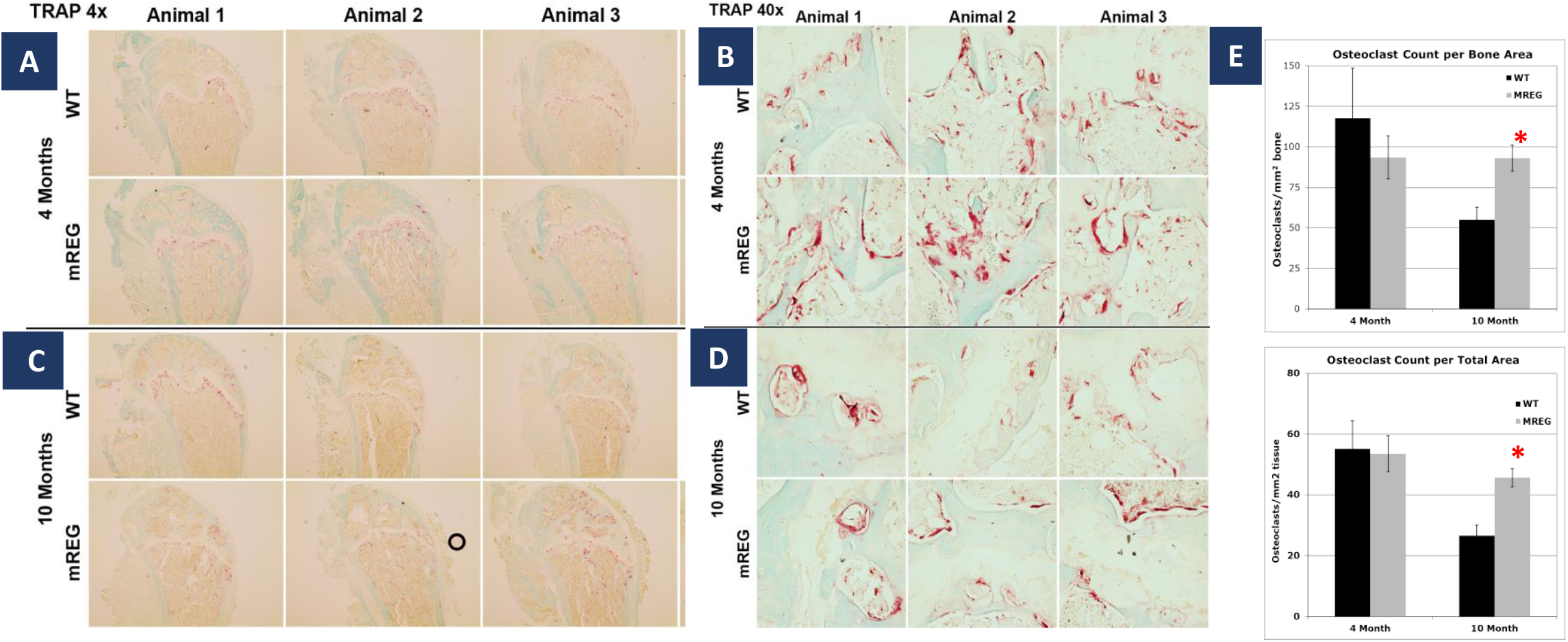
Osteoclast activity was assessed by TRAP staining in 4 and 10-month-old mice femurs (n=4-5). At 4 months, the osteoclast count was not different in WT vs. MREG mutant mice (A, B). At 10 months, differences were observed in the TRAP-stained cells for MREG mice (C, D). There was a significant increase in osteoclast numbers per total area in MREG mice at 10 months (E, * indicate statistical difference at p< 0.05).

### MREG^-/-^ bone marrow cells differentiation and function ex vivo

Primary bone marrow cells harvested from MREG^-/-^ and WT showed similar osteoclast differentiation over time (Fig. 4A). Immunoblotting did not show any difference for master osteoclast differentiation markers, such as NFATC1 and c-Fos (Fig. 4B). However, the resorption pit activity was significantly reduced in MREG mutant osteoclasts (Fig. 4C,D) (*p*< 0.05).

**Fig 4.**
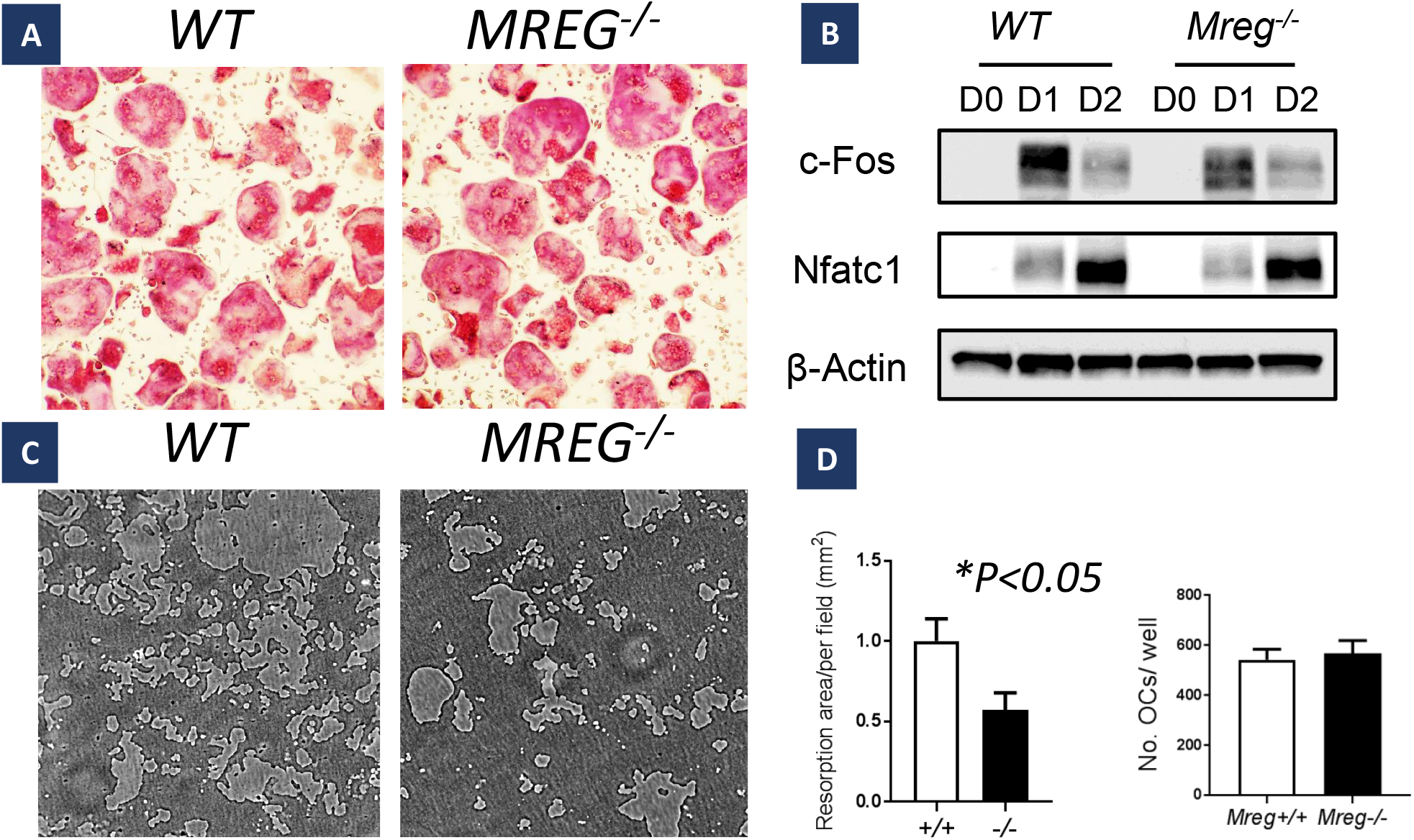
Bone marrow-derived osteoclasts from WT and MREG mutant (-/-) mice differentiate ex vivo similarly, as shown by TRAP staining (A) and Western Blot for osteodastogenesis master marker Nfatcl (B). Bone resorptive capacity is altered in MREG mutant (-/-) mice (C) with statistically significant differences in resorption pits generated by MREG mutatnt-derived osteoclasts (D) (p<0.05). Quantification of the resorptive area was statistically lower in MREG mutant mice.

### MREG^-/-^ mice showed higher total body lean mass

EchoMRI was used to assess the ratio of lean to adipose mass as reported (33). We observed a statistical difference between MREG^-/-^ and WT mice. The fat/bone ratio decreased for MREG^-/-^ at 10 months (Fig. 5A, B) (*p*< 0.05). The lean mass number was statistically higher for older MREG^-/-,^ although they did not differ in weight (Fig. 5A).

**Fig 5.**
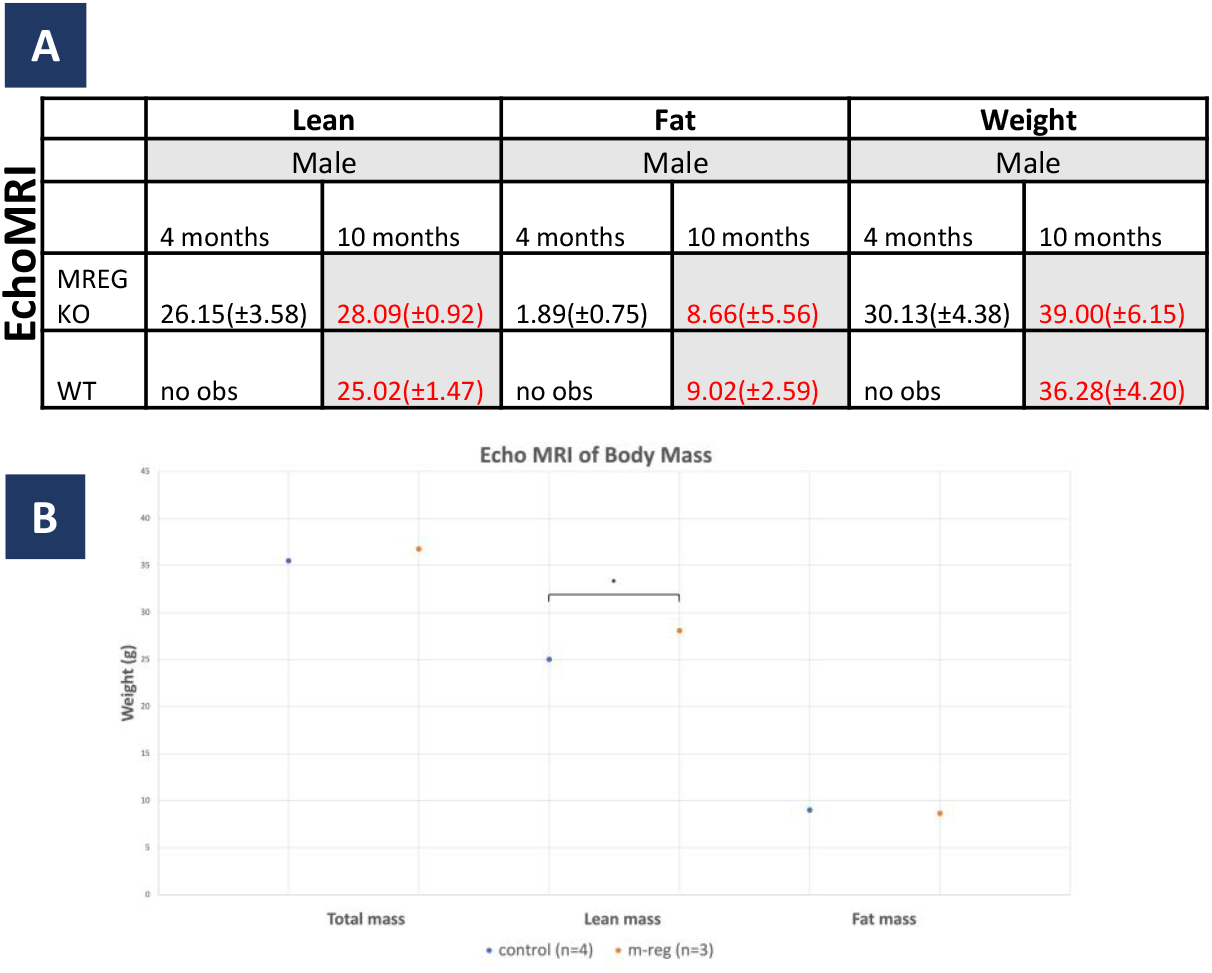
Quantitative analysis of lean and fat content of WT and MREG knock out (KO) mutant mice at 10 months of age via EchoMRI. Lean mass was significantly higher in MREG mutant male mice compared to WT at 10 months of age (A in red, B). WT mice at 4 months were not evaluated (not observed = no obs).

## DISCUSSION

This study aimed to evaluate bone phenotypic changes and primary osteoclast effects of melanoregulin deficiency. Through imaging, histological and cell culture methods, this small study suggests an important function of MREG in bone and possibly adipose tissue accumulation. This study shows a contradictory role of MREG deficiency, where it positively affected bone maintenance as observed by computed tomography, but also increased osteoclast numbers as observed histologically. Further, *ex vivo* culture of MREG^-/-^ bone marrow-derived cells shows no difference in cell numbers as they differentiate, but a reduction in their resorptive capacity. The paradoxal observation of increased osteoclast numbers in vivo concurrent with bone-sparing phenotype suggests reduced per-cell resorptive function may exceed the compensatory increase in cell numbers. Potential mechanisms of impaired fusion or altered osteoclast lifespan, upregulation of RANKL or decreased OPG to maintain normal resorption is warranted (26, 34, 35).

It was found that MREG function impairment significantly favors bone density as the mice age while decreasing marrow adipose tissue in the growth plate. Fat accumulation in the marrow is a physiological but undesirable bone phenotype acquired with aging that can lead to frailty and limited regenerative capacity. The phenotype of WT compared to MREG mutants could also be clearly associated with other undesirable metabolic changes such as obesity. This is supported by the finding that the MREG^-/-^ mice are leaner compared to WT, as measured by EchoMRI at 10 months, despite higher bone mass and no statistical difference in weight. The systemic metabolic alterations in MREG^-/-^ mice, including the localized reduced adiposity in bone marrow and increased lean mass is curious but it is important to acknowledge that bone marrow adipose tissue represents less than 1-2% of total body fat (36). Further, MREG deficiency could indirectly affect bone through mechanical loading or hormonal mechanisms independent of direct MREG function in bone cells (37, 38). Further investigation of circulating factors and inflammatory mediators are recommended for these mice besides level of expression of MREG in various types of cells (including osteocytes and osteoblasts), conditional deletion and rescue experiments. In addition, if MREG selectively affects trabecular bone, this site-specific finding will require further investigation.

We have a tentative hypothesis that MREG plays a significant role in cell fate of BMSCs. Since MREG has been shown to be inversely correlated with AKT-mtor signaling, it is possible that its absence will increase mtor and downstream beta-catenin signaling leading to increased osteogenesis and decreased adipogenesis (39-41). Further, autophagy regulation has been associated with metabolic disorders, thus, it is conceivable that MREG disfunction may affect overall systemic metabolism(42). Nonetheless, despite MREG’s documented role in autophagy (21, 43), we found no alterations in LC3 in MREG^-/-^ bone cells (not shown) for this cohort of mice, suggesting the bone phenotype may involve non-canonical autophagy pathways or autophagy-independent MREG functions.

True age-dependent studies to measure skeletal maturity, intermediate points and elderly time points in mice where age-related bone loss is pronounced is also needed to correlate MREG function with bone aging (44). It is a plausible hypothesis that MREG deletion could simultaneously enhance osteoblastogenesis while suppressing adipogenesis and affecting osteoclastogenesis (39, 45).

Our study calls for further definition of which cells are the critical direct target of MREG function in differentiation and explore the mechanism by which the marrow adipose tissue is reduced, and bone density is preserved. Further aging of mice is underway to assess later stages of bone phenotype. After a more thorough characterization of the phenotype of the mice with data on adipose tissue area, fat-related markers, metabolic markers, cell lineage regulation, extensive bone characterization (including sex as a biological variable) and monitoring of autophagy, it will be possible to more holistically assess MREG roles (46, 47). As autophagy regulates cellular energy, as well as aminoacid, glucose and lipid metabolism, the loss of autophagy homeostasis in adipose tissue and other tissues can have drastic effects on local and global metabolism (48, 49).

## CONCLUSION

MREG deficiency in male mice seems to affect osteoclast numbers *in vivo* and function and favor lean mass preservation over fat accumulation in bones and body composition as mice mature/age. This study will evolve into a thorough investigation seeking a deeper understanding of MREG and autophagy in bone and systemic metabolism. Understanding MREG function can provide future therapeutic modalities in inflammatory and metabolic diseases possibly related to autophagy.

## Acknowledgements and funding

EchoMRI was supported by P30DK056350 awarded to the Nutrition and Obesity Research Center at the University of North Carolina at Chapel Hill. mCT studies were kindly supported by Eric Livingston and Ted Bateman at the School of Biomedical Engineering at UNC-CH. Other mice experiments were supported by UNC Pilot Funds (to P.M.) and NIH R01 DE022465-01A1 (to K.B.B)

## Author contribution

**M.M:** Investigation, Formal Analysis, Validation, Writing-Reviewing and Editing. **V.Dp**.: Investigation, Validation, Writing – First draft. **S.T**.: Investigation, Formal Analysis, Writing – Reviewing and Editing. **S.W.W**.: Investigation, Validation, Writing – Reviewing and Editing. **K.B**.: Methodology, Resources, Writing – Reviewing and Editing, **P.M**.: Conceptualization, Methodology, Resources, Supervision, Project administration, Visualization, Writing – Original draft, Reviewing and Editing.

